# Simulated docking reveals putative channels for the transport of long-chain fatty acids in *Vibrio cholerae*

**DOI:** 10.1101/2022.01.26.477967

**Authors:** Andrew Turgeson, David Giles, Bradley Harris

## Abstract

Fatty acids (FA) play an important role in biological functions, such as membrane homeostasis, metabolism, and as signaling molecules. FadL is the only known protein that uptakes long-chain fatty acids in Gram-negative bacteria, and this uptake has traditionally been thought to be limited to fatty acids up to 18 carbon atoms in length. Recently however, it was found *Vibrio cholerae* has the ability to uptake fatty acids greater than 18 carbon atoms and this uptake corresponds to higher bacterial survivability. Using *E. coli’s* FadL as a template, *V. cholerae* FadL homologs *vc1042, vc1043*, and *vca0862* have been folded, simulated on an atomistic level using molecular dynamics, and analyzed revealing the FadL transport channels. For *vc1042* and *vc1043* these transport channels have more structural accommodations for the many rigid unsaturated bonds of long-chain polyunsaturated fatty acids, while *vca0862* was found to lack transport channels within the signature beta barrel of FadL proteins.

**IMPORTANCE:** Fatty acids are important precursors for membrane phospholipids as well as energy sources for bacteria. *V. cholerae’s* uptake of long-chain fatty acids has not been studied to the atomistic level at this point, and by doing so, the resulting putative pathways show important structural details of the transport protein. Not only do the transport channels have unique differences between one another, the predicted transport mechanics of *V. cholerae’s* protein homologs differ substantially from the *E. coli* FadL’s pathway found from X-ray crystallography studies.

## INTRODUCTION

In Gram-negative bacteria the transport of exogenous long-chain fatty acids (LCFA) across the outer membrane leaflet is mediated by FadL (1, 2, 3). FadL’s ability to acquire LCFAs grants versatility in carbon source utilization, providing a selective advantage for survival. In the case of *Vibrio cholerae*, the causative agent of cholera, this bacterial robustness may have ecological and medical implications. In this paper we will discuss the importance of bacterial FA synthesis and uptake. Then, we will compare novel structural models of *V. cholerae* FadL homologs with that of *E. coli’s*, highlighting both conservation and divergence in the proteins.

Fatty acids (FA) are molecules with a carboxylic acid head group and an aliphatic tail group of varying length and saturation. FAs are used primarily as building blocks for cell membranes, but also supply energy, and can be used as signaling molecules (4). Fatty acids can be acquired from exogenous sources as well as being synthesized *de novo*. However, many organisms (such as *Homo sapiens*) require specific exogenous sources of FAs for specific metabolic functions (4). In humans, this can be immune system regulation, blood clotting, neurotransmitter biosynthesis, cholesterol metabolism, and phospholipids for the brain and the retina (5).

In nature, plants typically have a limited synthesizing capacity that produces polyunsaturated fatty acids (PUFA) up to only 18 carbons (many plants are still capable of monounsaturated and unsaturated FAs for waxes and seed storage lipids) (6). However, plants are generally the only producers of n-3 (ω-3) and n-6 (ω-6) where the first unsaturated carbon starts on the 3rd or 6th carbon from the tail methyl group. Oddly, there are some heterotrophic bacteria (*Vibrio* and *Pseudomonas*) that can also produce the typically plant based n-3 PUFAs (7). Mammalian cells possess cytoplasmic fatty acid synthase (FAS) a major producer of 16-18 carbon atoms (which are also the most common cellular FAs in mammals) (6). Typically, plants and animals do not create the higher order (> 20 carbons) unsaturated fatty acids; instead, these longer chain FAs are commonly produced by marine protists and microalgae (6, 7, 8). It is widely known that fish, mollusks, and crustaceans tend to have high concentrations of the longer chain FAs such as eicosapentaenoic acid (EPA, 20:5) and docosahexaenoic acid (DHA, 22:6) (5). It is thought that all PUFA in food webs originate from primary producers, where organisms further up the food chain have only the ability to modify the FA by bioconversion and elongation as they pass through the food web (i.e., trophic upgrading) (4). Thus fish, mollusks, and crustaceans which have a diet of microalga and protist have higher concentrations of the longer chain PUFAs, but have a lessened ability for FA conversion to long PUFAs than freshwater fish (4).

In bacteria, FAs are primarily used as components for the phospholipid bilayer of the membrane. These membrane phospholipids are constantly being synthesized, modified, recycled, and degraded to maintain membrane homeostasis and to respond to environmental stressors (9, 10). Free FAs are released during these processes, constituting important sources of metabolic energy (9). Fatty acid biosynthesis involves a stepwise carbon elongation and unsaturation until the FA is of appropriate length and unsaturation. Further maintenance of membrane dynamics can be mediated by enzymes acting on constructed phospholipids, such as desaturases, cis/trans isomerases, and cyclopropane synthases (11).

Fatty acid synthesis pathways are highly conserved between bacteria and eukaryotes, the differences being the resulting fatty acids synthesized by bacteria tend to be slightly shorter, generally lack poly-unsaturation, and the monoenoic C18 acids have different double bond positions (12). Bacteria use type II fatty acid synthesis (FASII), which starts with an acetyl-CoA carboxylase complex (ACC) interacts with a biotin-dependent enzyme catalyzing an irreversible carboxylation of acetyl-CoA to produce malonyl-CoA. The resulting malonyl-CoA is used for the elongation cycle which extends the growing fatty acid with consecutive reduction, dehydration, reduction and condensation reactions by various fatty acid biosynthesis (Fab) enzymes (13, 6).

The elongation of FAs is costly, and as a result, all bacteria characterized to date have the capacity to uptake exogenous FAs (14). The pathway for long-chain FA uptake in Gram-negative bacteria begins with the transmembrane protein FadL (2, 15) to transport the FA into to periplasmic space, where it is then delivered through the inner membrane to FadD (acyl-CoA synthase or fatty acid-CoA ligase). FadD uses adenosine triphosphate (ATP) along with a FA, producing adenosine monophosphate (AMP) and P_2_ O_7_^-4^ (PPi) and a FA bonded to a coenzyme A (CoA) (16). The FA-CoA can then be shortened in beta oxidation producing a shorter FA tail and generating energy. Alternatively, a FA-CoA is the first component of a FAS elongation cycle, should the specific needs of the cell require a longer chain FA.

As previously stated, it was believed that enteric bacteria, such as *E. coli* and *V. cholerae*, were only able to acquire up to 18 carbon length FAs (17). However, over the past decade several Gram-negative pathogens have been shown to assimilate and respond to exogenous PUFAs (18, 19, 20, 21). In the case of *V. cholerae*, this increased uptake is likely due to its natural ecosystem of tropical climates where marine algae and protist are the bases of the aquatic food web. The uptake of PUFAs allows the incorporation of these long-chain FA into the cell envelope, and this incorporation has been shown to affect the membrane permeability, motility, biofilm formation, and antimicrobial resistance of the bacterium (22). Bile, along with mucus, in the human intestines has a large concentration of long-chain FAs in the form of phosphatidylcholine (23). Consequentially, *V. cholerae* has a lysophospholipase protein (VolA *vca0863*) capable of cleaving phosphatidylcholine and liberating fatty acids for bacterial uptake (24).

With increasing attention towards FAs and their effects in biology, the study of a species that exhibits broader capacity for the uptake and use of FAs presents an opportunity for comparison and elucidation of the uptake dynamics of the transmembrane protein FadL. In this paper we study the structure and functions of several *V. cholerae* FadL homologs (*v1042, vc1043*, and *vca0862*) using molecular dynamics and perform comparisons to the *E. coli* (*b2344*) FadL homolog.

## RESULTS

### Docking showed viability between the crystal structure and simulated results

There are no X-ray or NMR generated structure of the *Vibrio cholerae* FadL homologs, consequentially to find the structures of *vc1042, vc1043*, and *vca0862*, the protein sequences were folded using I-TASSER to find the tertiary structures. The resulting structures were simulated with the molecular dynamic (MD) simulator NAMD along with the known structure of *E. coli’s* FadL *b2344* henceforth referred as *b2344*. Various conformations of the equilibrated structures were docked with AutoDock using an array of fatty acids shown in Figure 1. The procedure is discussed in detail in the Materials and Methods section. To test the viability of the docking results, the FA library was docked with the original *E. coli* FadL *b2344* crystal structure (1T16) comparing the bindings of LDAO and C8E4 with the proximal bindings of the native LDAO and C8E4 molecules attached to the 1T16 PDB crystal structure. Figure 2 examines a frame of the resulting dockings showing a preference of the AutoDock bindings sites to primarily be the locations that were bound experimentally (25). However, the selected docking of the S3 kink binding site (residues highlighted in green) contains C8E4 molecules where in the original crystal structure C8E4 molecules were restricted to the low affinity binding site in the L3 and L4 extracellular loops. This may be due to the methodology of van den Berg, where LDAO and C8E4 competed for binding during the protein washing phase (25), while in simulated docking there was no binding competition.

**FIG 1.**
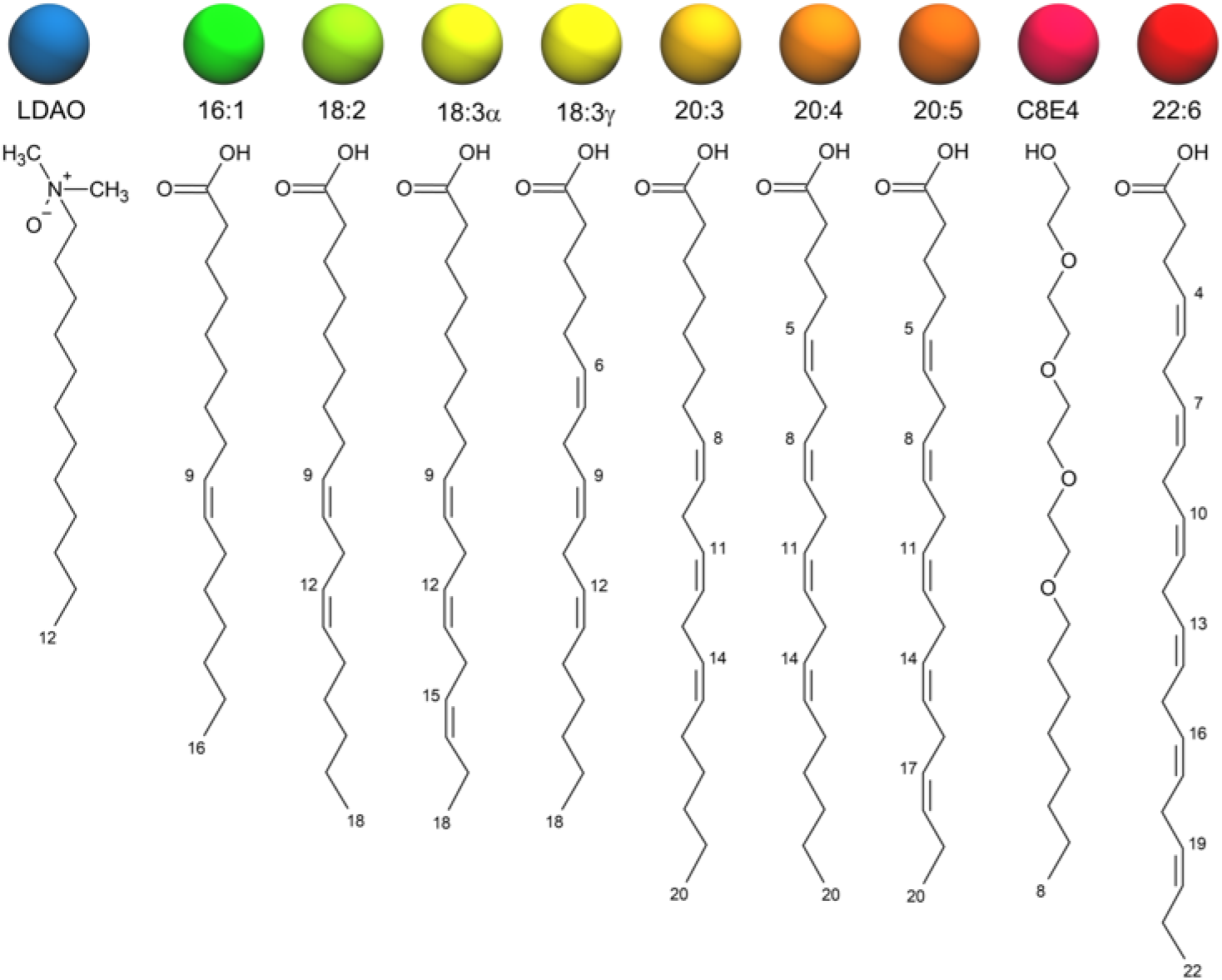
Fatty acids and detergents (LDAO and C8E4) used in docking the FadL homologs. The colored spheres show the color scheme associated with the corresponding fatty acid for images used throughout the paper.

**FIG 2.**
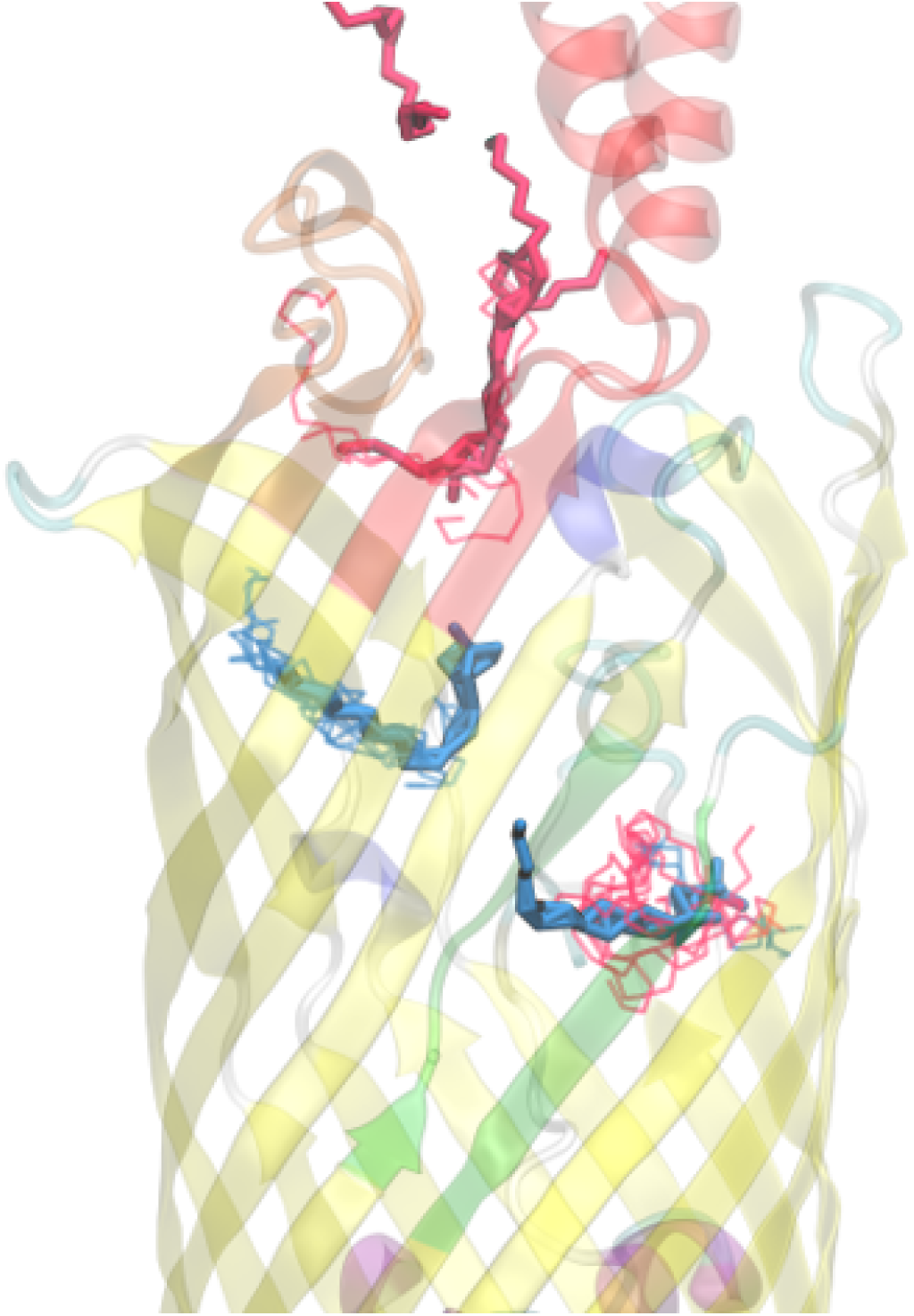
Docking of 1T16 with LDAO and C8E4 (lines) compared with the original LDAO and C8E4 (solid). While this is only one frame, the LDAO tends to be strongly correlated to the High affinity binding site while the C8E4 tends remain in the low affinity binding site or it bypass the transport channels and appears in the S3 kink (this occurrence is likely an effect of AutoDock’s ligand placement algorithm).

### The mean shift algorithm shows node cluster loci

Figure 3 shows the original *b2344* 1T16 crystal structure with the native detergents outlining the low affinity binding site, the high affinity binding site, and the S3 kink (25). The AutoDock binding within the 1T16 crystal structure in conjunction with a cluster analysis using a mean shift algorithm found the location of the low affinity binding site (Node 1), the high affinity binding site (Node 2), and the S3 kink (Node 3) by using the clustering of the docked FAs.

**FIG 3.**
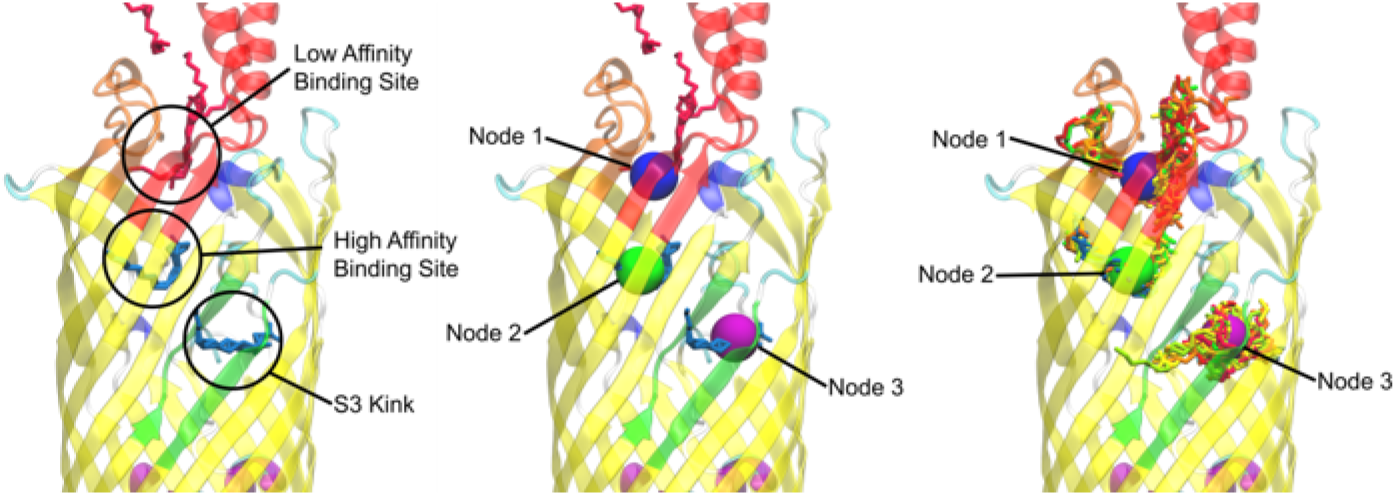
Mean shift based nodal analysis of the 1T16 docking. (Left) the original detergents and locations of the low affinity binding site, the high affinity binding site, and the S3 kink. (Center) The location of the nodes found by the mean shift algorithm. (Right) A frame of the FA clusters from docking that the mean shift algorithm used to generate the nodes. (Bottom) Color coding of each FA tested to show clustering by type.

The cluster analysis was also performed for each of the FadL and *V. cholerae* and the equilibrated *b2344* homolog structures, resulting in the nodal locations seen in Figure 4. The nodes from equilibrated *b2344*, Figure 4A, shows almost identical nodal locations as the X-ray 1T16 structure Figure 3. A small difference between the high affinity binding site where Node 2 tended to be closer to the center of the FadL beta barrel in the equilibrated structure). Exception aside, this demonstrates the NAMD equilibration *b2344* FadL structure tended to retain important structures during the simulation.

**FIG 4.**
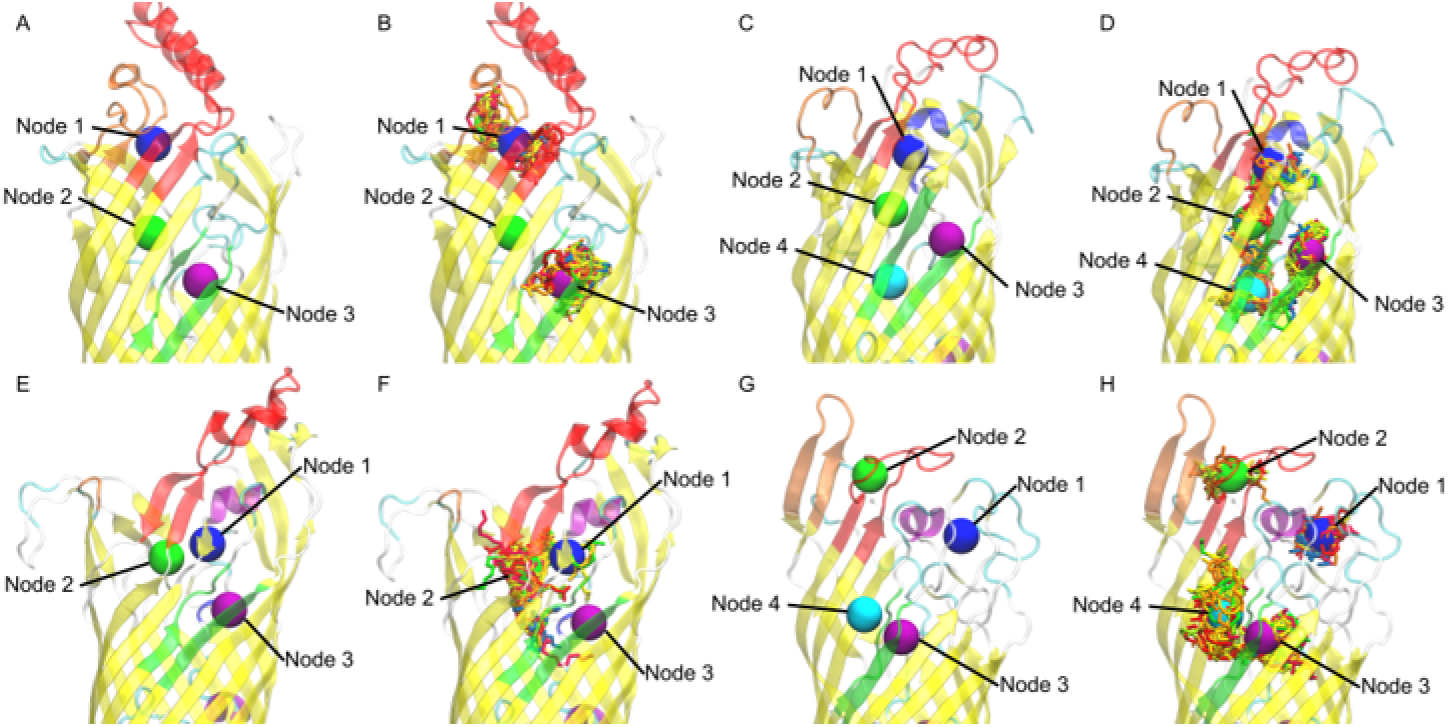
Nodal Locations of (A) *E. coli*, (C) *vc1042*, (E) *vc1043*, (G) *vca862*. Example frame of FA clusters versus mean shift generated nodes for (B) *E. coli*, (D) *vc1042*, (F) *vc1043*, (H) *vca862*.

### The nodal analysis shows preference for certain node loci

The FAs were categorized based on proximity, with each docked FA being prescribed one node per frame. With 10 FAs per type per frame and 50 frames, 5,000 FAs were assigned to each FadL homolog - giving a reasonable statistical model. The resulting docked locations were summarized in Figure 5.

**FIG 5.**
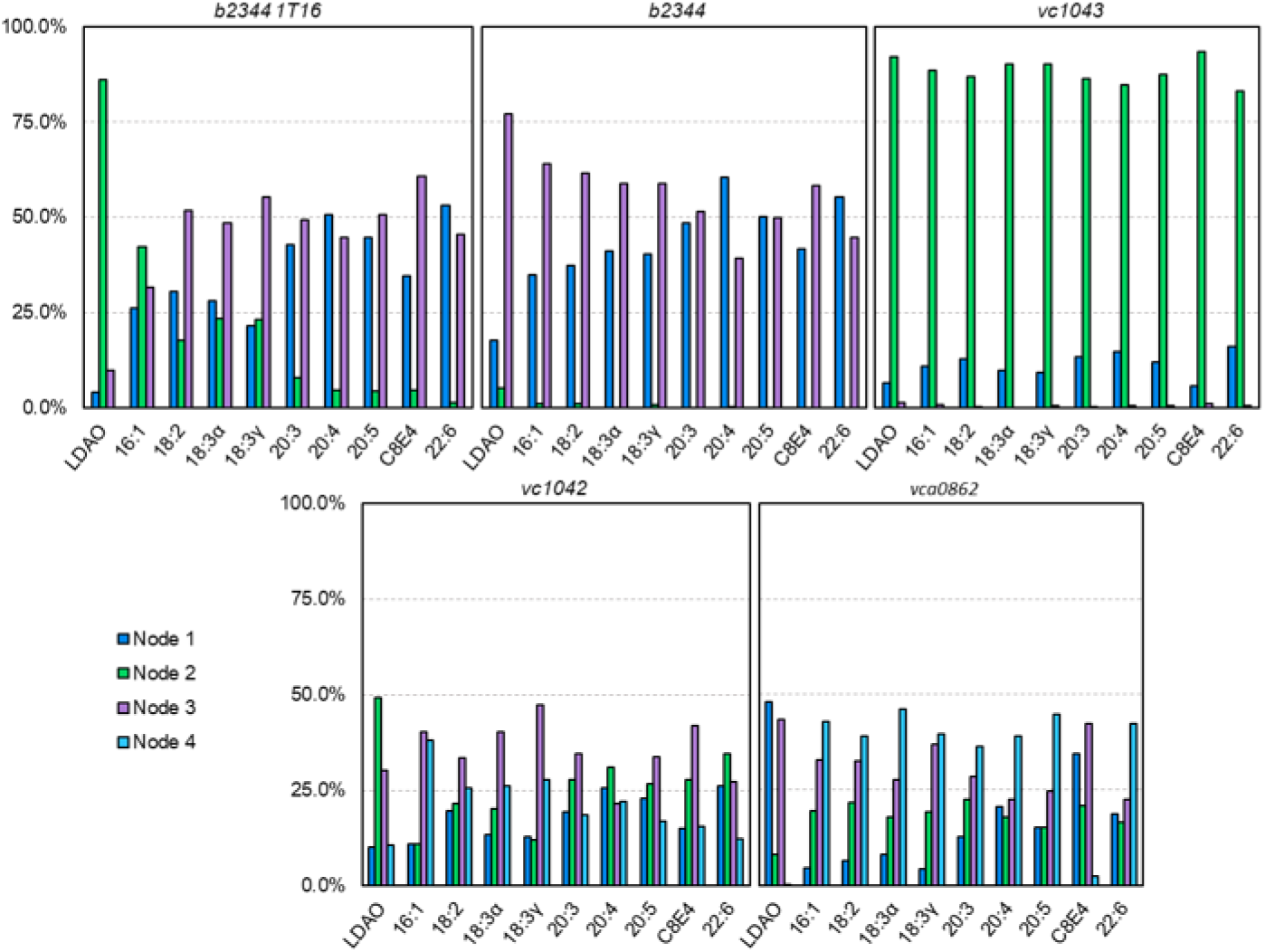
Charts of FAs by type located around certain nodes. The % is out of the 500 docked instances of each FA type over the 50 frame trajectory. The *b2334, b2334*, and *vc1043* docking resulted in three nodes, while *vc1042* and *vca0862* resulted in four nodes.

For the 1T16 test case, LDAO had a strong affinity for the high affinity binding pocket (Node 2) with 86.0% of the LDAO molecules docked appearing in or around the high affinity binding site. The other small molecules such as 16:1, 18:2, 18:3α, and 18:3γ also showed clustering in the Node 2 region (42.2%, 17.8%, 23.4%, and 23.2% respectively). This is reasonable due to the original 1T16 structure having a LDAO molecule bound to the high affinity binding site (25) (the Node 2 locus), where other similarly shorter chained FAs could also fit into the open pocket. Interestingly, the S3 kink (Node 3) tended to have more docked FAs than the low affinity binding site (Node 1) which may be due to the tubular cavity of the S3 kink region, providing more surface area for FAs to bind to than the more open low affinity binding site. Amongst all the FAs tested, the average docking binding energy for Nodes 1 and 3 of 1T16 were -8.975 and -8.976 kcal/mol respectively, indicating a very close average binding energy. Examining 18:2 in specific, the binding energy of 18:2 with Node 1 was better (−9.29 versus -8.97 kcal/mol), but AutoDock propagated more 18:2 FAs on Node 3. This is likely due to the AutoDock algorithm finding it more difficult to dock the Node 1 area due to a smaller binding channel, even if the binding channel has a better binding energy. Another example of this phenomena is 22:6 having the best overall binding energy when it was found in the high affinity binding site (−12.65 versus -9.73 and - 10.11 kcal/mol), however, this only occurred with 1.4% of 22:6 dockings because the high affinity binding site was originally bound to LDAO – a shorter chain FA.

The equilibrated *b2344* docking revealed that the high affinity binding site had fewer dockings than the other sites. This indicates that due to the vacancy of FAs during simulation, that the high affinity binding site was smaller and did not dock many FAs. Further investigation revealed that small positional changes in the high affinity binding pocket residues - particularly ALA153, ILE155, and LEU200 impeded the binding pocket channel, and greatly reducing the ability for FAs to fit in the binding pocket. Unlike the high affinity binding site, the S3 kink did appear to have substantial binding, indicating that there is not a conformational change in the S3 binding pocket during FA transport, but instead a shift in the gated channel between the high affinity site and S3 kink as proposed by van den Berg (25). The size and saturation of the FA did have an effect on the docking. Typically, the longer the FA carbon chains and more unsaturated, the affinity for Node 1 was increased and the affinity for Node 3 was decreased - mirroring the 1T16 dockings.

The *vc1042* docking revealed a visible main channel. It is predicted that the FAs move from Node 1 to Node 2 to Node 4 and then to Node 3, the S3 kink (Figure 4D). The clustering of FAs did not show much preference for any one of the four nodes with the exception of Node 3, where the totaled percent FAs located at Nodes 1, 2, 3, and 4 were 17.5%, 26.2%, 35.0%, and 21.3% respectively. As expected of a *V. cholerae* homolog there was no discernable difference in FA tail length or saturation which is reasonable due to *V. cholerae’s* ability to uptake long-chain fatty acids.

In *vc1043*’s docking, Nodes 1 and 2 were in close proximity to one another as seen in Figure 4F, the major difference between the two being Node 2 is the locus of the high affinity binding site in the *E. coli* homolog. *vc1043* showed a strong favoritism for Node 2 with a total of 88.3% of all FAs appearing in the Node 2 region. Figure 4F illustrates the FA’s tendency to funnel around Node 2. Only 0.6% of FAs were found in the S3 kink region (Node 3), alluding to a conformational mechanism to allow passage of the FA.

The docking of *vca0862* showed a high affinity for the outer portion of the S3 kink (Node 4). This is unexpected based on the premise that FAs travel through the beta barrel in *E. coli*. Node 4 does tend to have a more pronounced indention making docking more ideal than some other locations; however, the docking did not factor in the LPS which encompassed the outer perimeter of the FadL beta barrel, which would leave little room for FAs. Autodock’s current atom limit prevents a system with LPS included. These results indicate that the *vca0862* beta barrel was not in an open conformation for the 50 frames used for docking and suggests that there is a very large conformational change that possibly starts with FA binding to Nodes 1 or 2 in the extracellular loop region. It is important to note that the lower C-score in the I-TASSER folding, -1.08, may have a part in this resulting barrel structure, however, additional foldings and molecular dynamic runs using different programs (data not shown) were performed to test the viability of alternatively folded *vca0862* sequences; none of which had any notable differences in the structure.

### Eqiulibrated *E. coli b2344* fatty acids show similar channels as the X-ray crystal structure 1T16

To determine any important residues in the transport of FAs, the docked homologs were searched for any residues within 3 Å of each of the docked FAs. These residues were agglomerated, and each residue found was counted for recurrences. The resulting table (Table 1) shows the twenty residues that were found to interact with the docked FAs most often.

**TABLE 1.**
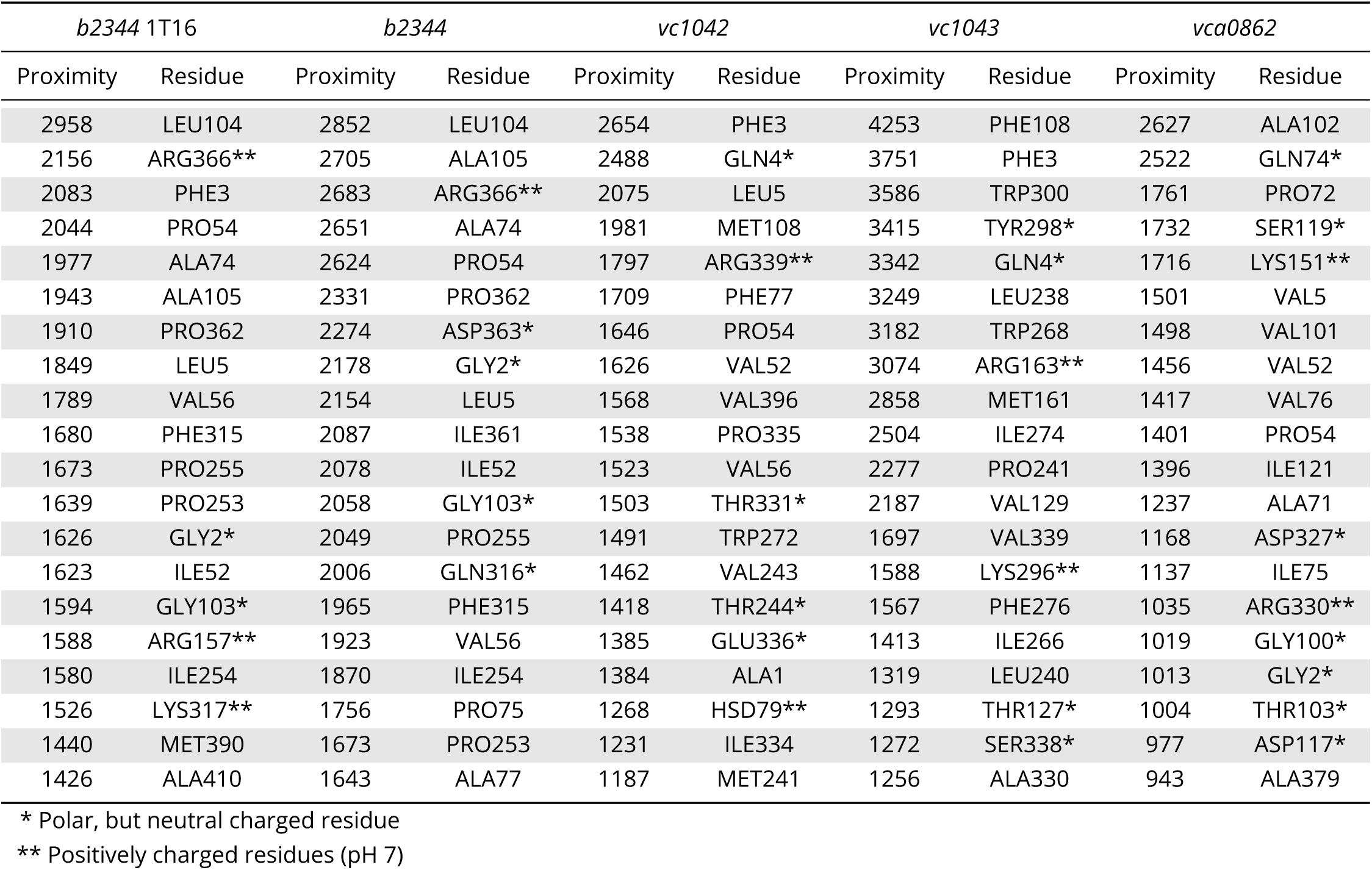
Residue count for residues found within a 3 Åproximity of each FA for each frame. For each FadL homolog 100 FAs were docked for each of the 50 frames, giving a maximum residue count of 5,000.

For the *E. coli b2344* homolog dockings, the residues found most frequently were those of the low affinity binding site and the S3 kink. This was expected, as the nodal analysis determined that the majority of FAs were docked in the Node 1 and 3 regions. While not in the same proportions, many of the same residues were found for both the *b2344* 1T16 and the equilibrated *b2344* structures. Residues PRO253, ILE254, PRO255, and PHE315 have a reoccurring presence in the low affinity binding sites for both structures Figure 6A and C. Residues GLY2, LEU5, PRO54, VAL56, ALA74, GLY103, LEU104, ALA105, PRO362, and ARG366 are commonly found in the S3 kink region. The majority of these residues are nonpolar except for the polar glycines GLY2 and GLY103 and the positively charged arginine, ARG366. The arginine headgroup faces towards the S3 kink pocket indicating an affinity for carboxyl groups of FAs which is confirmed by the number of FA carboxyl headgroups in the proximity of ARG366 during docking. This could indicate an orientation of FA with the tail group facing the outlet before egress of FA through the S3 kink. The RMSD for the heavy atoms of these residues tends to be between 1.1Åand 1.7Å, alluding to a stable S3 kink structure even with the difference of a bonded LDAO in the S3 kink of 1T16.

**FIG 6.**
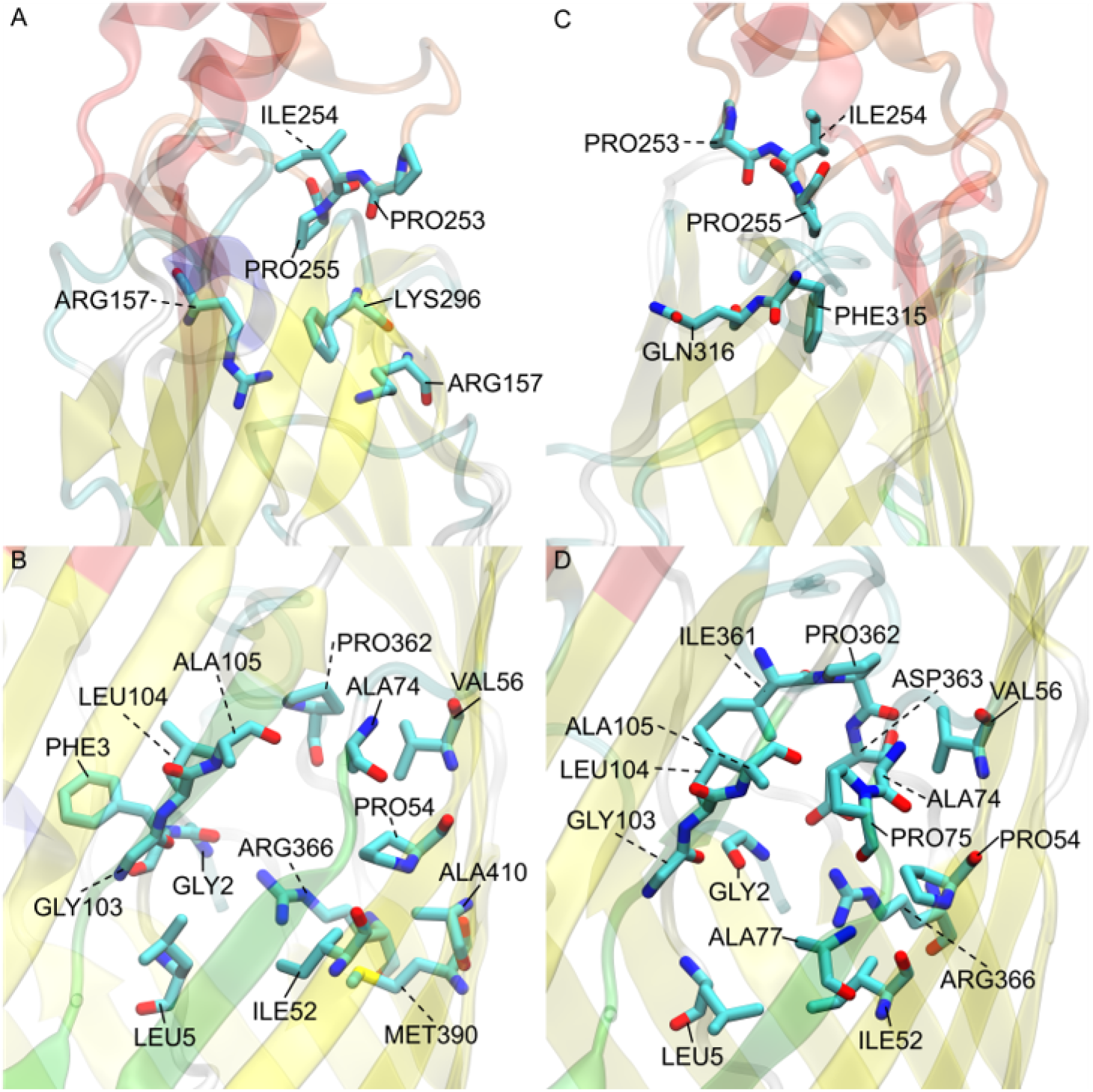
*E. coli* FadL binding residues of (A) 1T16 low affinity binding site residues, (B) 1T16 S3 kink, (C) *E. coli* low affinity binding site, and (D) *E. coli* S3 kink. The L3 loops are colored red, the L4 loops are colored orange, and the S3 kinks are colored green for reference. Perspective angles differ for easier observation of residues.

### *V. cholerae vc1042* docking generates a well defined transport channel

The *V. cholerae* homolog *vc1042* residues were primarily centered around the predicted transport channel Figure 7B. This channel tends to start from between the 5th and 6th extracellular loops (Figure S1B). The FA is expected bypass the N-terminal hatch (residues 1 through 5), and then past the N-terminal hatch through the S3 kink opening. The N-terminal hatch in the docking does not restrict transport as predicted in the *E. coli* homolog *b2344*. This is somewhat unexpected as, generally, a FA transport protein would have some selection mechanism specific to FAs. The residues that line the channel are primarily hydrophobic, with a few exceptions GLN4, HSD79, THR244, THR331, GLU336, and ARG399 which are all hydrophilic (ARG339 also having a positive charge). These residues are placed periodically throughout the channel in such a way that it could be the FA headgroup’s attraction to these residues that guide the movement of the FA through the channel in a specific orientation. The channel seems to end at the S3 kink as with the *E. coli* homolog. A similar experiment with the docking of the bottom half of the homologs showed that there was a discontinuity from the main channel to any docking channels found in the bottom of the protein reinforcing the hypothesis that the S3 kink is the FA egress point. Oddly, the channel shares a similar overlap of the *E. coli* homolog’s high affinity binding site location and N-terminal hatch domain, but interestingly the *vc1042* pathway bypasses the predicted hatch domain pathway used in *E. coli*. It is interesting that the original pathway (through the high affinity binding site location, and then through a tunnel created by a conformational mechanism occurring with the N-terminal hatch domain) may still exist somehow in the *V. cholerae* homolog. Whether or not this N-terminal hatch pathway is vestigial or is functional has yet to be determined at this time.

**FIG 7.**
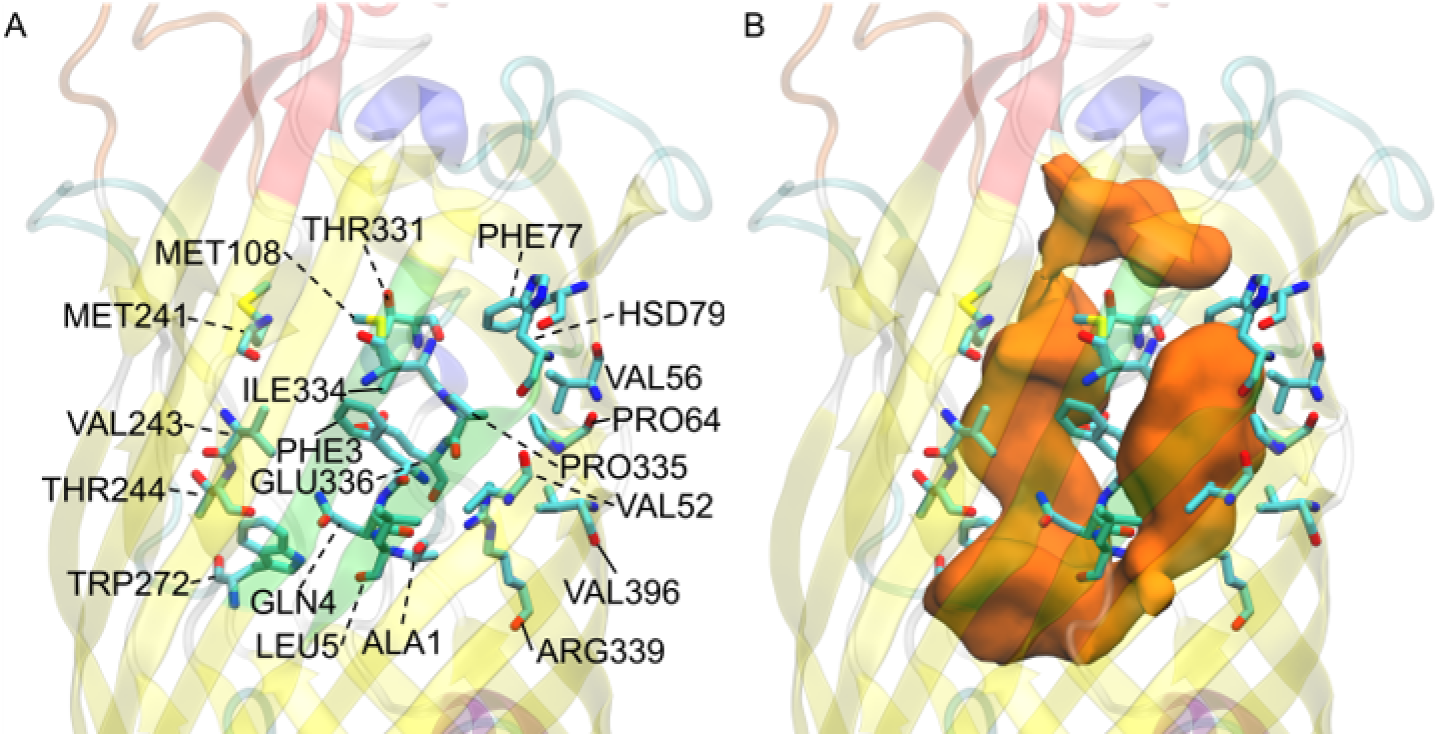
*V. cholerae vc1042* FA interacting residues; (A) residue names and locations, (B) predicted transport channel (orange) shown with its relation to the displayed residues. The L3 loops is colored red, the L4 loop is colored orange, and the S3 kink is colored green for reference.

### *V. cholerae vc1043* transport channel has a discontinuity revealing multiple predicted pathways

*V. cholerae* homolog *vc1043* has a very large flat channel that seems to funnel FAs to the N-terminal hatch domain (Figure 8). The entry of the channel can be seen in the supplemental material (Figure S1C) where the undefined low affinity binding region is again found between the base of the extracellular loops. The channel leads to the N-terminal hatch opposite the S3 kink. The N-terminal hatch domain rests in the same position as the *E. coli* 1T16 structure and the channel overlaps the general area of the high affinity binding. This could indicate an evolutionary adaptation to combine the low affinity and the high affinity binding sites found in *b2344* favoring a more direct pathway, but leaving the mechanisms of the N-terminal hatch domain which would play the same role for the *vc1043* homolog as it does for the *E. coli* homolog. This would require the N-terminal domain to act as a hatch that opens and closes for FA transport, unfortunately this mechanism at the atomistic level has not been elucidated for accurate prediction. Alternatively, the *vc1043* channel spans further down than the transposed high affinity binding site ending below the N-terminal hatch and opposite the S3 kink. This could be an alternate pathway that follows the *vc1042* pathway, but with some selection mechanism to cross the remainder of the channel. Again, this proposed pathway has yet to be substantiated from experimentation. The docked FAs did not appear in the S3 kink pore, likely due to LYS130 from the fourth beta strand, S4, positioned parallel to the S3 kink that appears to be attracted to GLU50, SER106, and ASN107 as well as the backbone oxygens of the S3 kink residue GLY109. This attraction causes LYS130 to fill the S3 transport pore and prevent docking (and possibly FA transport). This could be a selection mechanism that may determine the resulting FA position or FA type. The channel of *vc1043* was found to be composed generally of hydrophobic residues. The exceptions to this are GLN4, THR127, TYR298, and SER338 which are hydrophilic, and ARG163 and LYS296 which are positively charged. Previously, it was postulated that the hydrophilic residues in *vc1042* guided the FAs through the channel through hyrdophilic residue interactions, but *vc1043* hydrophilic residues are restricted to the top of the conical channel. There- for there are no hydrophilic residues present in the vacant channel to guide the polar FA headgroup through the channel. It is predicted that the polar head groups bind to the hydrophilic residues at the top of the channel for alignment. Directional positioning of the FA is yet to be determined experimentally, but FA orientation may play an important role with a positively charged lysine (LYS130) residue blocking the S3 kink pore.

**FIG 8.**
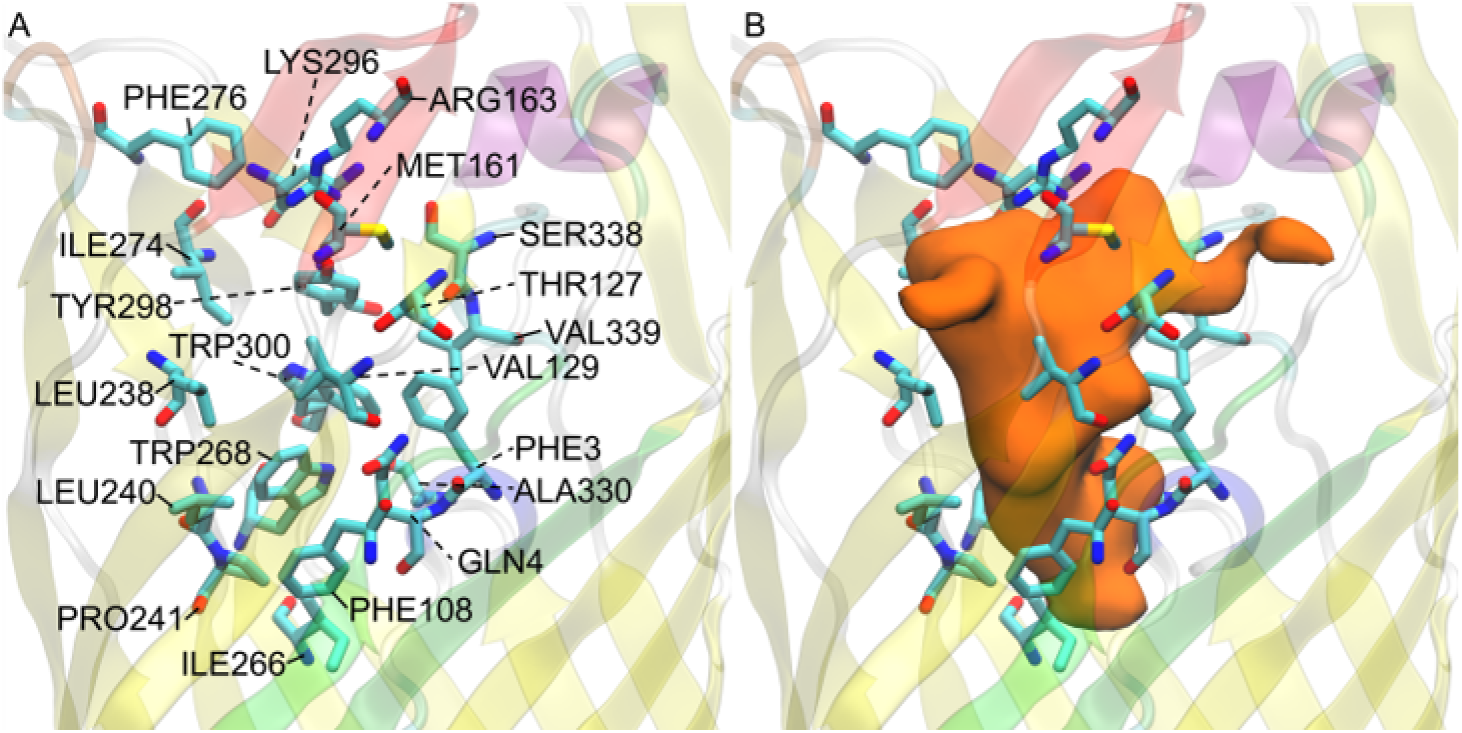
*V. cholerae vc1043* FA interacting residues; (A) residue names and locations, (B) predicted transport channel (orange) shown with its relation to the displayed residues. The L3 loops is colored red, the L4 loop is colored orange, and the S3 kink is colored green for reference.

### *V. cholerae vca0862* docking reveals a transport channel external to the beta barrel

The docking of *V. cholerae* homolog *vca0862* revealed that the majority of docking sites did not occur within the beta barrel structure of the FadL protein, but rather along the outer barrel primarily around the S3 kink (Figure 9B). This appears to be due to the substantial lack of open space for FAs to be docked on the inner portion of the beta barrel. Oddly, there seems to be a pathway from between the L3 and L4 loops that goes down the side of the protein and to the outside of the S3 kink as shown in the supplemental material (Figure S2). The results imply that any molecule of similar size to a FA would be able to make its way through the side channel unless there was some interplay with the interface of the LPS and lipid bilayer to create some sort of selectivity mechanism. The channel between the outer portion of the S3 kink and the predicted initial binding sites between the L3 and L4 extracellular loops tends to close off depending on the L3 and L4 conformations. These L3 and L4 conformations may be the selectivity mechanism that this homolog uses to ensure the uptake of FAs instead of bactericidal compounds. Many of the docked FA were found within the S3 kink, where the internal cavity of the S3 kink would be vestigial if the FAs are transported to the predicted egress point without entry of the FA into the FadL beta barrel structure. Unless the s3 kink has an orientation mechanism to help FAs diffuse passively through the membrane. This vestigial S3 kink cavity agrees with the secondary docking of the bottom portion of the protein, where no FA pathways were found from the S3 kink to the periplasmic end of the FadL protein.

**FIG 9.**
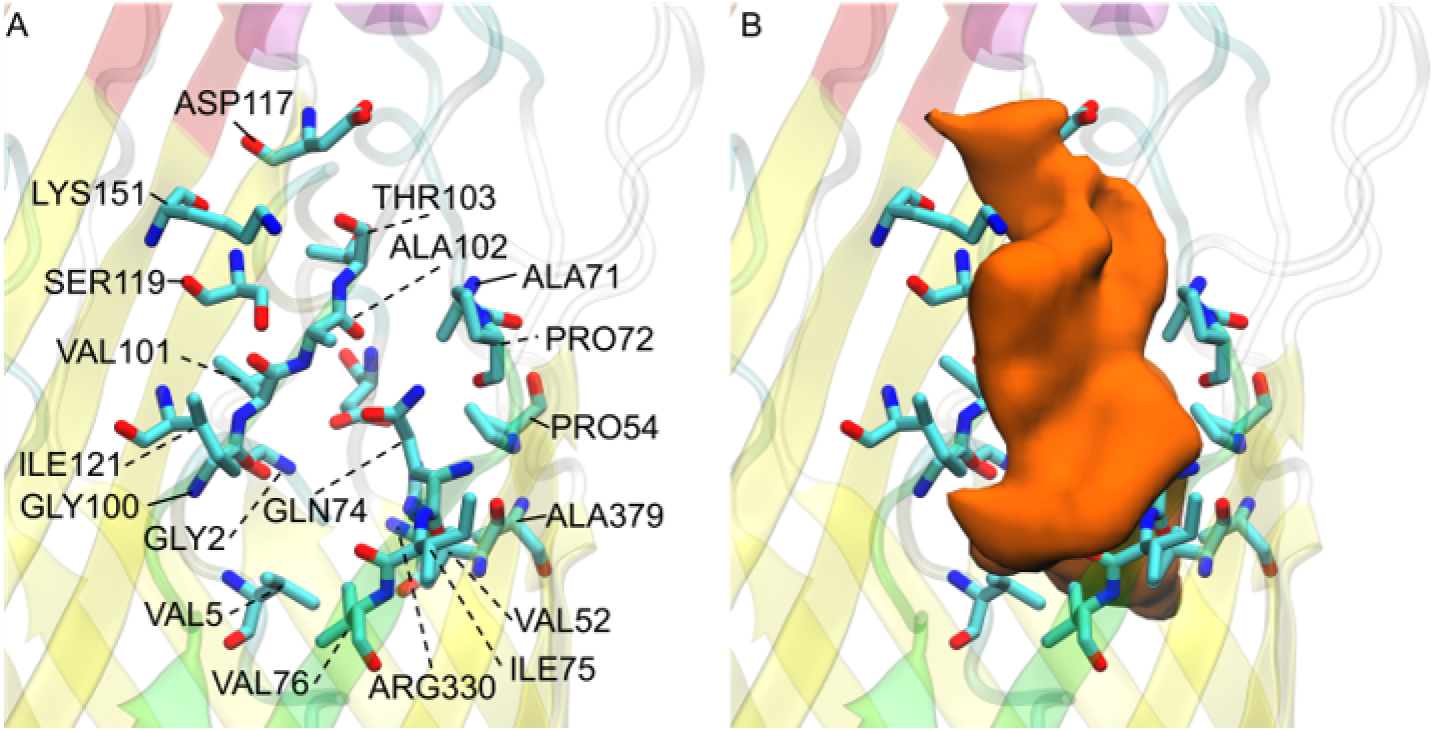
*V. cholerae vca0862* FA interacting residues; (A) residue names and locations, (B) observed transport channel (orange) shown with its relation to the displayed residues. The L3 loops is colored red, the L4 loop is colored orange, and the S3 kink is colored green for reference.

### Simulations and docking agree the S3 Kink is the fatty acid point of egress

To verify that the S3 kink is the egress point first suggested by Hearn *et al*. (26), the membrane layer location after equilibration was checked for the possibility of membrane diffusion. The resulting lipid bilayer headgroups or the polar heavy atoms of the LPS were shown in relation to the D3 kink pore (Figure 10). This pore was typically found at the upper portion of the LPS polar region indicating a strong affinity for the polar headgroups of the FA with the polar LPS residues, indicating a good possibility for assimilation into the LPS bilayer and passive diffusion into the periplasmic space. Additional docking studies (not shown) using the bottom portion of the FadL structures also revealed no such pathways through the lower portion of the N-terminus loop that fills the lower beta barrel for either *V. cholerae* or *E. coli*.

**FIG 10.**
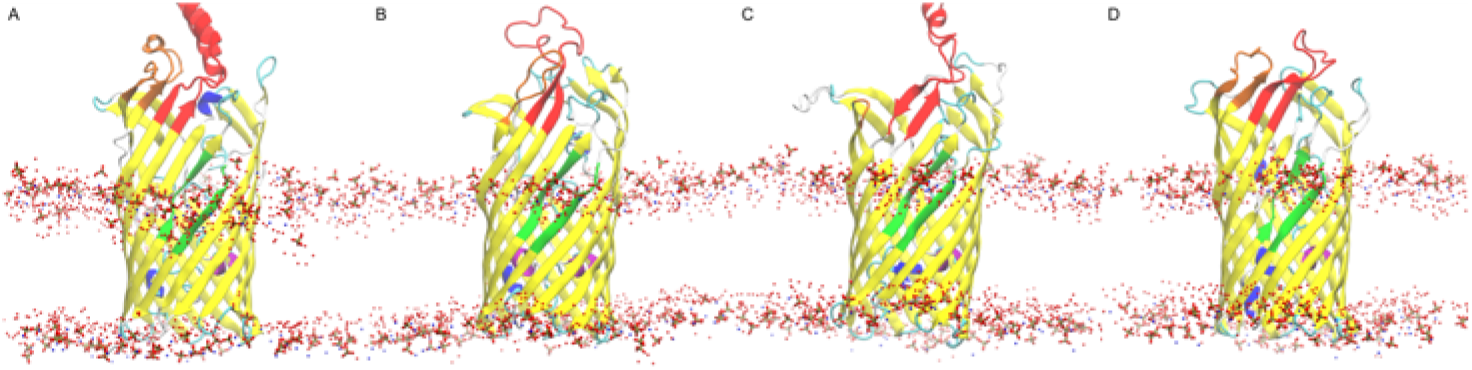
Lipid bilayer polar atom locations with respect to the (A) *b2344*, (B) *vc1042*, (C) *vc1043*, and (D) *vca0862* FadL proteins after equilibration and docking.

### Docking energies show that binding is stronger in the presence of fatty acids

10 docking conformations were produced per ligand creating a total of 100 docking conformations per FadL homolog frame. With 250 frames docked, 25,000 conformations were generated overall. The best conforming (lowest energy) are shown in Table S1. Similarly, the averaged docking energies by FA are given in Table S2.

Simulated docking results indicate that the original crystal structure *E. coli* 1T16 tended to have the most energetically favorable docking with respect to overall average as well as the best individual FA docking conformations. This is likely because the 1T16 structure was generated with the FadL protein bound with LDAO and C8E4 in the structure when the PDB was generated, giving it the specific conformation needed for strong binding. It is also apparent that the docking energies are more favorable for the longer chain FAs (with exception to C8E4 which has a total length of 21 heavy atoms). This result is likely due to the fact that longer FA chains provide more surface area for binding. However, many of the longer chained polyunsaturated FAs tend to have a hairpin tail due to the cis unsaturated portions. The uptake of these longer chained FAs would likely require some internal mechanisms for FA uptake that compensates for these rigid sections of the FAs, although these compensation mechanisms have not been found computationally.

## DISCUSSION

The atomistic study of FadL homologs reveals the structure and transport channels of *V. cholerae* homologs *vc1042, vc1043*, and *vca0862*. The *E. coli* controls showed agreeable results when compared the X-ray crystallography study (25). All homologs tended to share similarity in their low affinity binding site locus at the base of the L3 and L4 extracellular loops, but the structure of extracellular loops themselves tended to deviate from the original template. The reason for the deviations has has yet to be determined, but it may have an effect on nascent protein passage through the cell membrane, localization of the protein in the membrane with respect to the LPS, or as guide for FAs into to the transport channel.

The equilibration of *b2344* in comparison to the X-ray structure (1T16) shows that there are conformational shifts in the binding sites for FAs that are closed without the presence of FAs. *vc1043* shows this similarity, where it appears the presence of FA are required for the span between the observed channel and the S3 kink domain to activate and allow passage. The trajectories over 50 nanoseconds gives a wide range of conformations for the FA to propagate on the proteins, and for neither *b2344* or *vc1043* to reveal a contiguous channel through docking emphasizes the importance of the FA-protein interaction. This is not the case with *vc1042*, where the channel is fully expressed without FAs being present. The N-terminal hatch residues ALA1, GLY2, PHE3, and GLN4 were conserved throughout all of the homologs, as well as their tertiary structures and positions. These residues may play a part in conformational changes that allow selective passage of FAs through the cell (25, 26). However, the channels presented for the *V. cholerae* homolog *vc1042* suggests the possibility that this structure can be vestigial in the transport of FAs, but retain the conserved sequence as part of the signal peptide sequence (27), the protease recognizing the signal sequence ALA-GLY-PHE-GLN as part of the processing site. This is subject to the protease responsible for the cleavage, which is has not been discovered at this time.

The resulting structure of *vca0862* shows that the channel is on the exterior of the beta barrel. This may be due to the folded structure, but multiple foldings and equilibrations across different software did not predict any significant differences in the structure. Investigation using RefSeq (28) on FA uptake revealed that *V. cholerae* strains that have these homologs all contain a copy of *vc1042, vc1043*, (chromosome I) and *vca0862* (chromosome II) (data not shown). *vca0862* has not been shown in any studies to be the definitive protein responsible for FA transport, with many bioinformatic searches of FadL neglecting *vc1042* and *vc1043*. Additionally, in studies with gene expression of *V. cholerae* strain N16961 between *in vitro* and *in vivo*, the expression of *vca0862* was low compared to the other FadL sequences (29). The relatively low expression of *vca0862* and the lack of a suitable channel for FA transfer may indicate that this protein may be a result of a loss of function adaptation (30). Interestingly the data presented by Xu *et al*., reveals that expression of *vc1042* increases *in vivo* (in rabbits) in comparison to *in vitro* (growth in LB). The inverse was true for *vc1043* with a reduction in expression *in vivo*. The effect of FA concentration in the lumen as opposed to the FA devoid LB, may be a selection mechanism for *vc1042* which appears to have a larger, more complete, and less selective channel.

The atomistic structures of *Vibrio cholerae* FadL homologs were found and analyzed. The channels of these homologs bring to light the complex nature of biological systems and the diverse machinery that is used to adapt to environmental conditions. While each homolog has unique characteristics, the exact nature of each homolog is still unknown, and additional studies into these characteristics will shed light on the diversification and expanded uptake capacity for not only *Vibrio* species, but the growing list of Gram-negative bacteria demonstrating fatty acid utilization versatility.

## MATERIALS AND METHODS

### Selecting the *Vibrio cholerae* homologs

*E. coli* M1655 FadL (accession number: NP_416846) was used as input for a BLAST (31) homolog search against all available sequenced *V. cholerae* strains of the pathogenic O1 and O139 serogroups. The search algorithm settings were set at 100 max target sequences, short queries, an expect threshold of 10, a BLOSUM64 scoring matrix, with gap costs determined by Existence: 11 Extension: 1, and a Conditional composition score matrix adjustment. No filters or masks were used to analyze the results.

The resulting proteins were reduced to unique sequences and *vc1042, vc1043*, and *vca0862* (accession numbers: WP_000856207, WP_001061938, and WP_000966057 respectively) were selected based on prevalence.

### Generating the FadL tertiary structures

The selected *V. cholerae* sequences were removed of the predicted signal peptide sequences, folded using the I-TASSER (32) standalone version, and compared to the known crystal structure of *E. coli’s* FadL from the RCSB database (PDB ID: 1T16) (25). The C-score determined by I-TASSER (on a scale from -5 to 2) for *vc1042, vc1043*, and *vca0862* were 0.58, 0.70, and -1.08 respectively. The resulting structures can be seen in the supplementary material Figure S3.

### Generating the membrane system

The resulting FadL structures including the *E. coli b2344* 1T16 structure were then each placed into a membrane using CHARMMGUI Membrane Builder (33). The *V. cholerae* homologs’ membrane had an outer leaflet of *V. cholerae* type 1 Lipid A, Core A, and 15 O1 O-antigen units. The *E. coli* outer leaflet was composed of *E. coli* type 1 Lipid A, Core R1, and 3 *E. coli* O1 O-antigen units (with 5 sugars per O unit). Both types had an inner leaflet of 67% phosphatidylethanolamine (PE) and 33% phosphatidylglycerol (PG). The structure of each of these molecules can be seen in the supplementary material, Figure S2.

### Equilibrating the membrane systems

The resulting simulation constraints generated by the CHARMM-GUI were then used in conjunction with NAMD (34) and CHARMM36 force fields (35). During simulations, Langevin dynamics were used to maintain constant temperature (310 K) and pressure (1 atm). The simulations were sized as 80Å x 80Å x 140Å and a flexible cell boundary was chosen for an anisotropic membrane system. A cutoff of 12Å was used along with a particle mesh Ewald (36) for electrostatic interactions. All equilibrations used a timestep of 2 fs and nonbonded frequency and full electrostatics calculated at every step. Each of the four protein systems were equilibrated for a minimum of 250 ns. The RMSD of the equilibration run for each protein tested can be seen in Figure S3. A Ramachandran plot was made to detect conformational outliers and determine a Z-score using MolProbity (37) of the homolog structures at 0 and 201 ns to determine the differences between structure of the and the initial and membrane equilibrated structures. The outlier residues found were sparse both spatially and numerically in both instances, indicating that the structures were of good agreement. Additionally, the Z-scores were within acceptable tolerances, where typically the absolute value being less than 2 (Figure A5 in the appendix).

### The docking of the FadL proteins

50 frames of the equilibrated FadL trajectories were taken (one every nanosecond), and for each frame the protein was isolated and aligned. An array of 10 ligands (Table S3)were then docked to each of the frames taken in addition to 50 instances of the 1T16 crystal structure. The 80×80×120 AutoDockTools gridbox binding region (closer to a 40Å x 40Å x 60Å box) was restricted to the upper extracellular region of the FadL proteins encasing the majority of the FadL proteins. To maintain the cis structures of the FAs, the unsaturated double bonds of the ligands were kept rigid during docking. Each docking used a genetic algorithm with a population size of 150, a maximum number of evaluations of 2,500,000, and a maximum of 27,000 generations.

## SUPPLEMENTAL MATERIAL

**TABLE S1.**
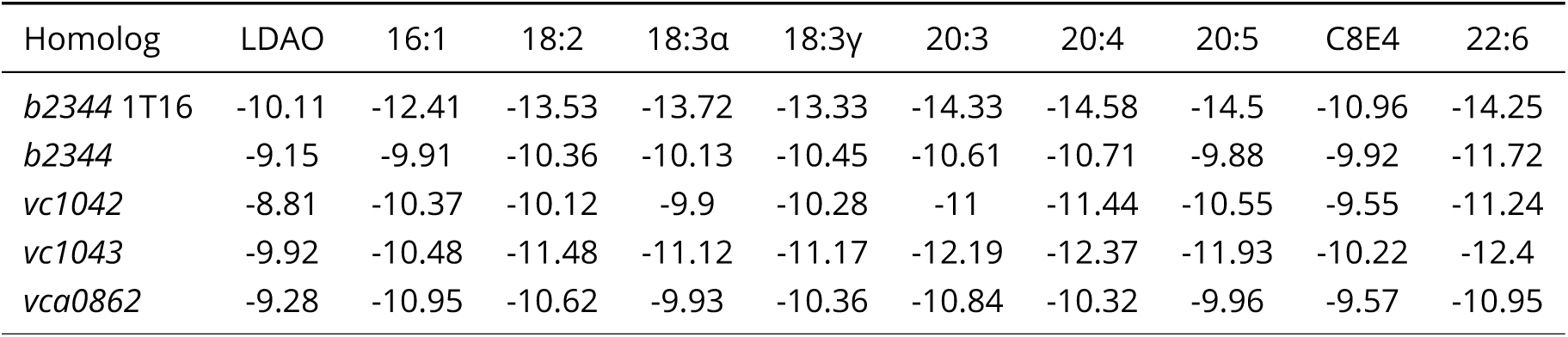
Best Conformation Energies from Docking (energy units in kcal/mol)

**TABLE S2.**
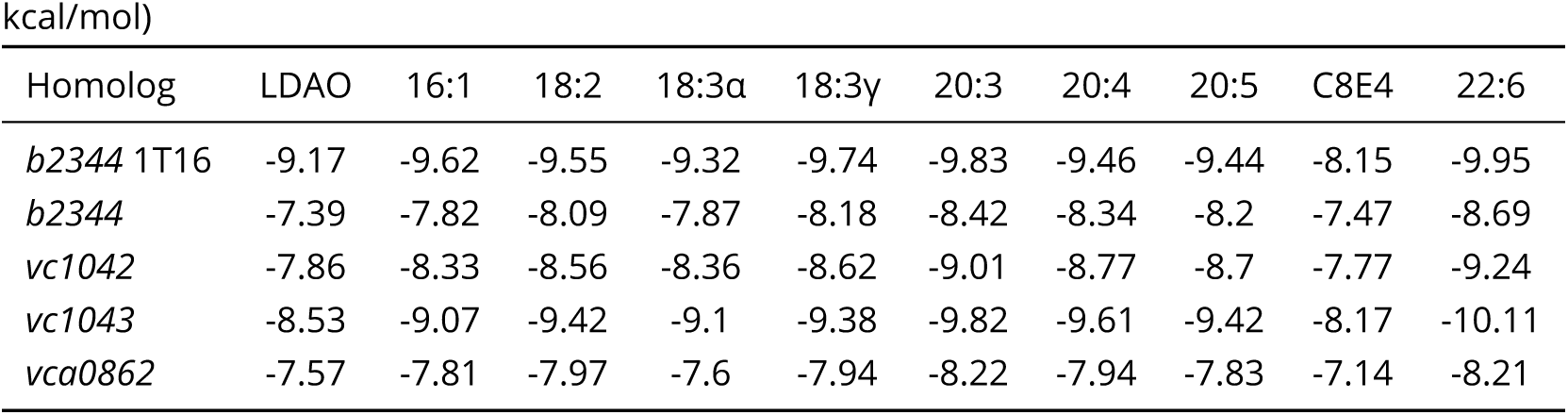
Overall Average Conformation Energy from Docking (energy units in kcal/mol)

| Homolog | LDAO | 16:1 | 18:2 | 18:3α | 18:3γ | 20:3 | 20:4 | 20:5 | C8E4 | 22:6 |
| --- | --- | --- | --- | --- | --- | --- | --- | --- | --- | --- |
| *b2344* 1T16 | -9.17 | -9.62 | -9.55 | -9.32 | -9.74 | -9.83 | -9.46 | -9.44 | -8.15 | -9.95 |
| *b2344* | -7.39 | -7.82 | -8.09 | -7.87 | -8.18 | -8.42 | -8.34 | -8.2 | -7.47 | -8.69 |
| *vc1042* | -7.86 | -8.33 | -8.56 | -8.36 | -8.62 | -9.01 | -8.77 | -8.7 | -7.77 | -9.24 |
| *vc1043* | -8.53 | -9.07 | -9.42 | -9.1 | -9.38 | -9.82 | -9.61 | -9.42 | -8.17 | -10.11 |
| *vca0862* | -7.57 | -7.81 | -7.97 | -7.6 | -7.94 | -8.22 | -7.94 | -7.83 | -7.14 | -8.21 |

**TABLE S3.**
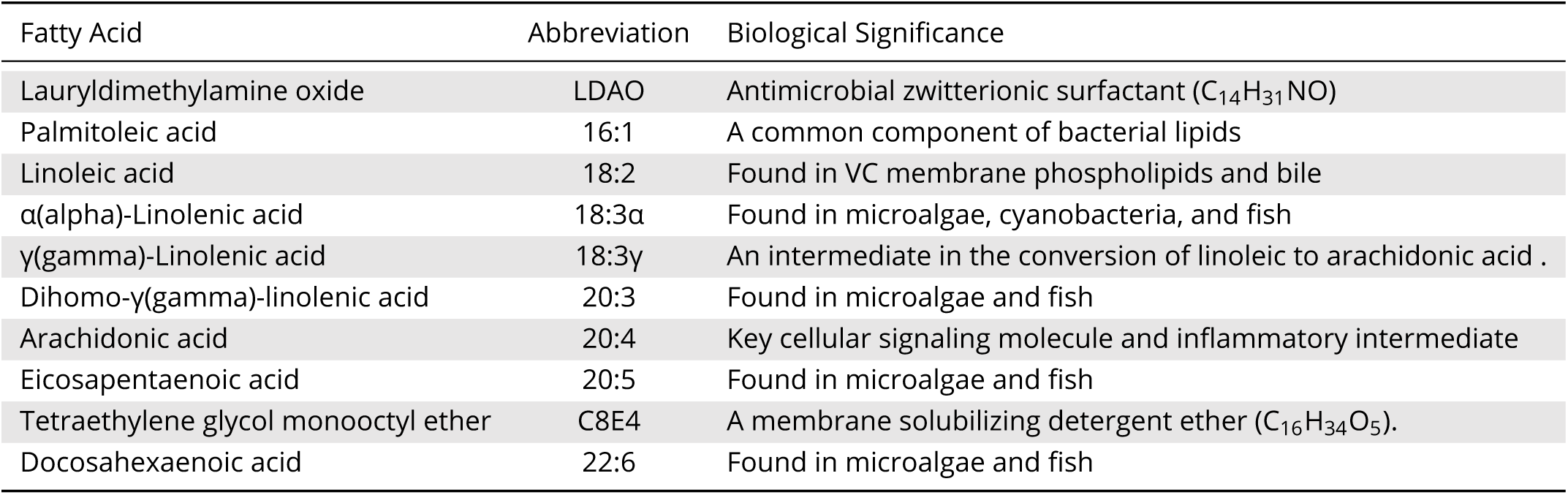
Fatty acids tested during docking

**FIG S1.**
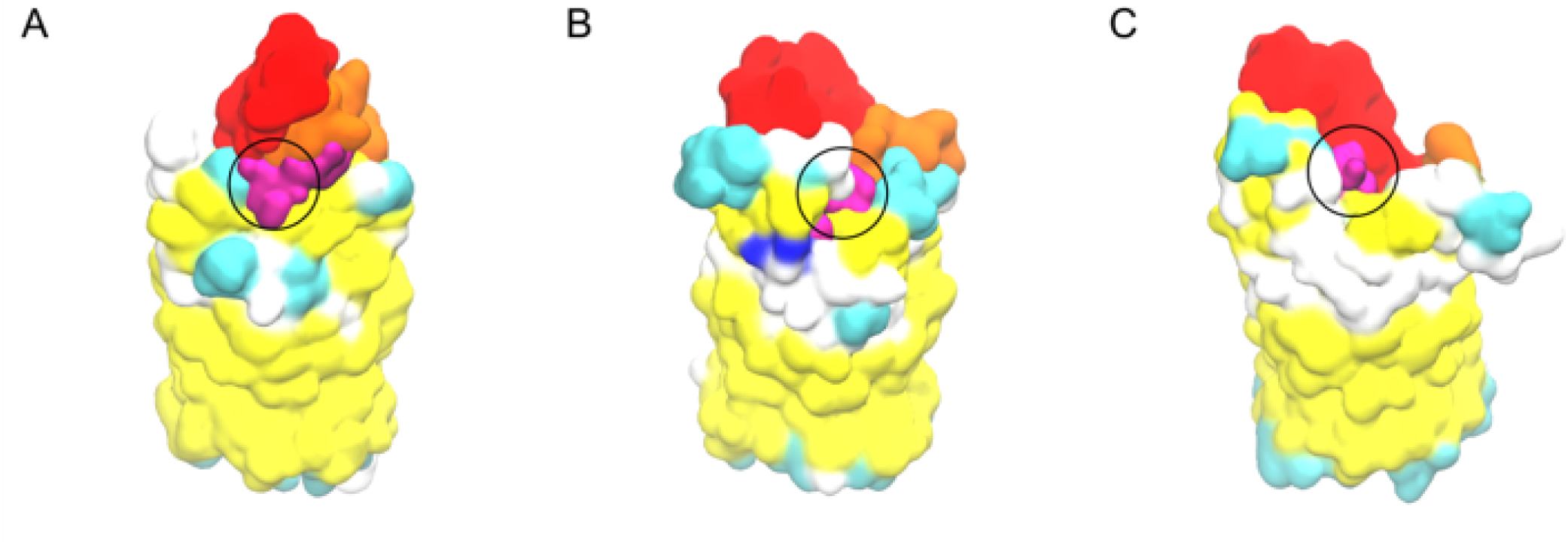
Surface view of FA transport channel entrance points (shown in purple) for (A) *b2344*, (B) *vc1042*, (C) *vc1043* FadL proteins after equilibration.

**FIG S2.**
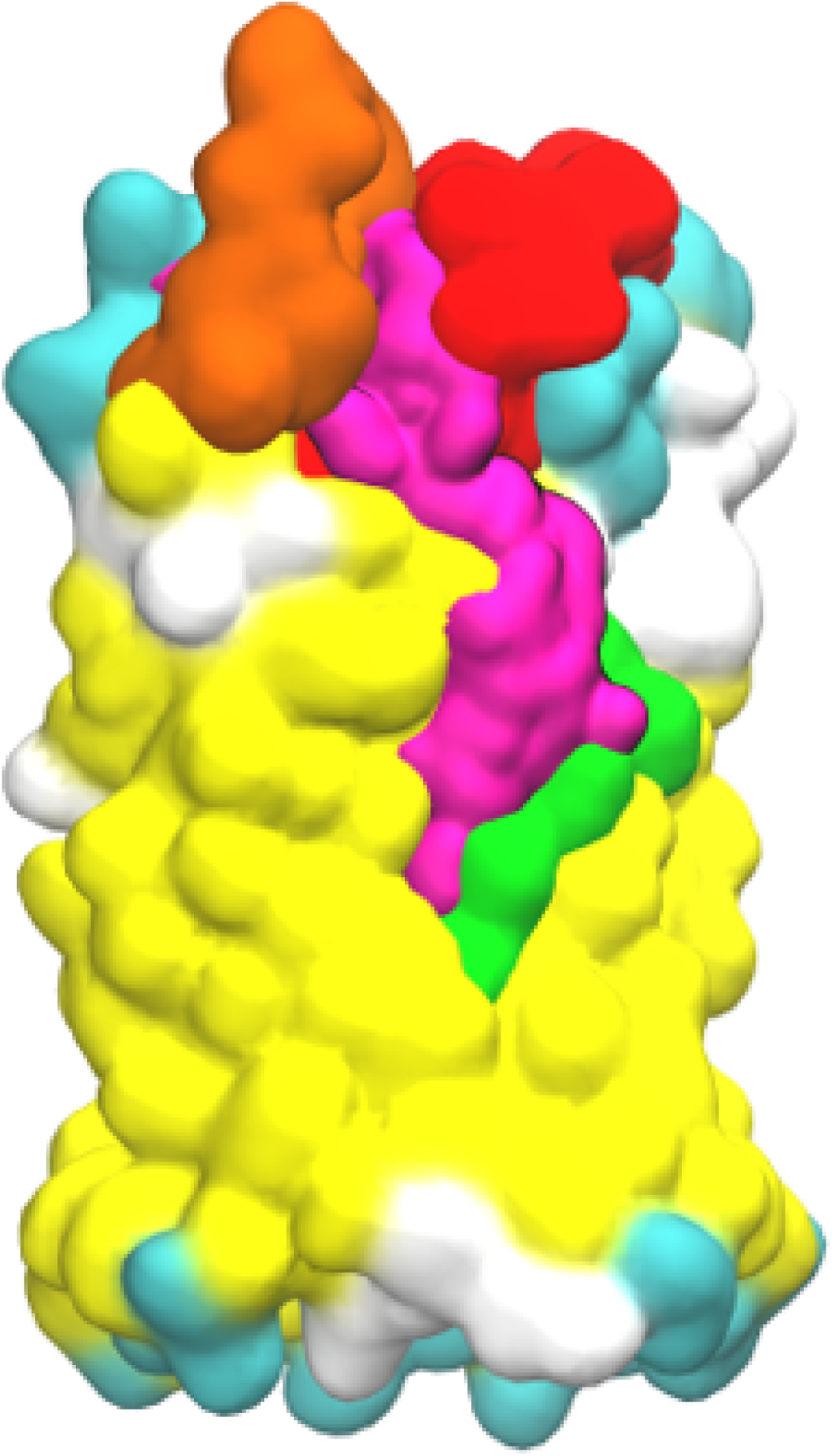
Surface view of FA exterior pathway (shown in purple) for *vca0862* FadL protein after equilibration.

**FIG S3.**
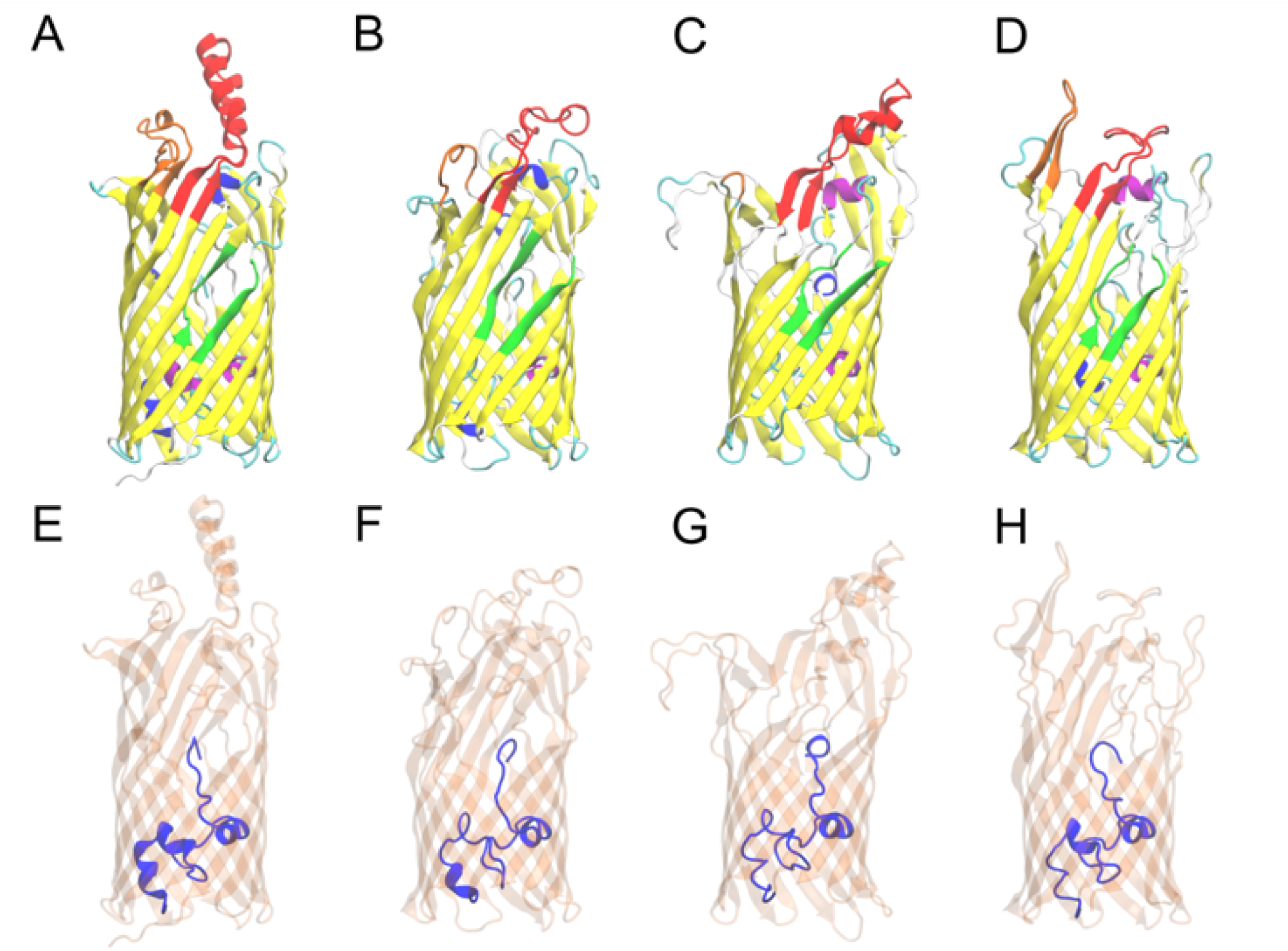
Comparison of I-TASSER folded structures. (A) the *E. coli b2344* 1T16 crystal strcuture compared to (B) *vc1042* (C), *vc1043*, and (D) *vca0862*. L3 and L4 loops are colored red and orange respectively, and the S3 kink is green for reference. (E-H) are views of the N-terminal hatch domain (blue) for *E. coli b2344, vc1042, vc1043*, and *vca0862* respectively.

